# Sonic Kayaks: Environmental monitoring and experimental music by citizens

**DOI:** 10.1101/167833

**Authors:** Amber GF Griffiths, Kirsty M Kemp, Kaffe Matthews, Joanne K Garrett, David J Griffiths

## Abstract

The Sonic Kayak is a musical instrument with which to investigate nature, developed during open hacklab events. Kayaks rigged with underwater environmental sensors allow paddlers to hear real-time water temperature sonifications and underwater sounds, generating live music from the marine world. Sensor data is also logged every second with GPS, time and date, allowing fine scale mapping of water temperatures and underwater noise that was previously unattainable using standard research equipment. The system provides the paddler with an extra dimension of senses with which to explore the underwater climate, while enabling citizens to gather data for scientific research. The system can be used as a citizen-science data-collection device, research equipment for professional scientists, or a sound-art installation in its own right, and has been implemented in a public setting at the British Science Festival 2016, demonstrating the considerable advantages of adopting transdisciplinary approaches during project development. Here we present instructions for building the open-hardware and open-source software, tests of the sensors used, and preliminary data demonstrating applications for the Sonic Kayak in marine climate and noise-pollution research.

## Introduction

The essence of scientific research is the exploration of the unknown and the discovery of something new. Likewise, understanding and explaining the world is one of the core foundations of the arts and music. With the benefits brought by the internet and distributed manufacturing for sharing protocols and hardware design, the power to do scientific research no longer needs to be confined to expensive research labs. With more accessible hardware comes the opportunity for more ambitious and adventurous science education and engagement that is inherently merged with the process of scientific research design and data collection. Combining the approaches of the sciences and arts provides opportunities for mutual benefits that are more than the sum of their parts.

Audification and sonification can be used to sense, explore, and understand data, and are essentially the auditory equivalent of data visualisation. Broadly speaking, audification is data sonification at its purest – simply transforming data directly into sound - while sonification is a broader term referring to conveying information using non-speech sounds (Hermann et al. 2011). Detecting changes in rhythmic patterns, pitch, volume, tempo, timbre etc. can give insights into changes in scientific model outputs or empirical data. The transformation of data directly into sound has proven its worth in a number of scientifically-linked applications, including Geiger counters which transform ionizing radiation levels into audible clicks, and pulse oximeters used in medical settings which emit higher pitches for higher blood oxygen concentrations. Audification is particularly suited to situations where you need to have access to a constant stream of data and be able to detect changes in that data while continuing with other activities. This is because the human ear can’t physically shut out sound as the eye can shut out visual information.

The Sonic Kayak project emerged from the Bicrophonic Research Institute (BRI), established by Kaffe Matthews and David Griffiths in 2014. Through ten years of international projects the BRI developed the Sonic Bike whose music changes depending on where the cyclist goes. Sound maps are made through residential research and workshop collaborations with producers and communities, and loaded onto the system on the bikes. As the cyclist pedals around the city, a GPS receiver on the bikes detects where they are and triggers site-specific sounds, played through a pair of bike-mounted speakers. Not an app and free of the internet, the Sonic Bike creates an outdoor listening experience for all, the antithesis of headphones - the cyclist becoming a performer and the passers-by the audience. Politics, time, cycling possibilities, architecture and finances bare on these new works. Sonic cycling hubs have appeared worldwide, in Finland, Houston, London, Brussels, and Berlin.

The Sonic Kayak project builds directly on the Sonic Bikes - we modified the open-source technology for use on boats and added underwater sensors to take a pure arts project into the realm of citizen-science. Underwater sounds, detected by hydrophones (underwater microphones), are played through speakers, along with data from two digital thermometers sonified in real-time. This allows the paddler to explore the underwater environment through sound. As with the Sonic Bike system, location-specific sounds can also be loaded onto maps and triggered in different places. Meanwhile, the sensor data is recorded with time, date, and GPS location, providing research-quality data of use for marine noise pollution and climate research. The Sonic Kayaks were launched at the British Science Festival (Swansea, 6-9 September 2016).

In this paper we outline instructions for building the open-hardware and open-source software for making a Sonic Kayak. We demonstrate the suitability of the sensors used and provide preliminary data from the hydrophones and digital thermometers, considering the potential applications in marine noise pollution and climate research. We discuss the approach of using open hacklabs for project development and the use of public events for data collection.

## Methods

### Sonic Kayak Hardware

#### Temperature Sensor Tests

For the Sonic Kayak system to work, we needed temperature sensors that could do two things: (i) Gather underwater microclimate data that is high enough quality for scientific research purposes, and (ii) Provide a live-feed of data for sonification purposes.

The standard temperature sensor for marine research is the TidbiT. These are often attached to marine buoys for long-term temperature data collection. The sensor is small (about the size of a bottle cap), there is a ‘shuttle’ to connect the sensor to a computer, and proprietary software to programme the sensor before deployment and download/graph the data after deployment. In total this system costs approximately £240. Because TidbiTs can not provide a live data feed, they were not suitable for the Sonic Kayaks, however we used a TidbiT (HOBO TidbiT v2 Water Temperature Data Logger) to check the data obtained from low-cost (£3) waterproof 1-wire digital thermometers (part number <u>DS18B20</u>).

A low-cost digital thermometer was wired up to a Raspberry Pi 2 computer (see Fig. 1 showing the adaptor which came with the sensor, and how to wire the adaptor into the Raspberry Pi) and code was written to read/write the sensor data each second (available open source here: https://github.com/sonicbikes/sonic-kayaks/blob/master/sensors/therm.py).

**Fig 1.**
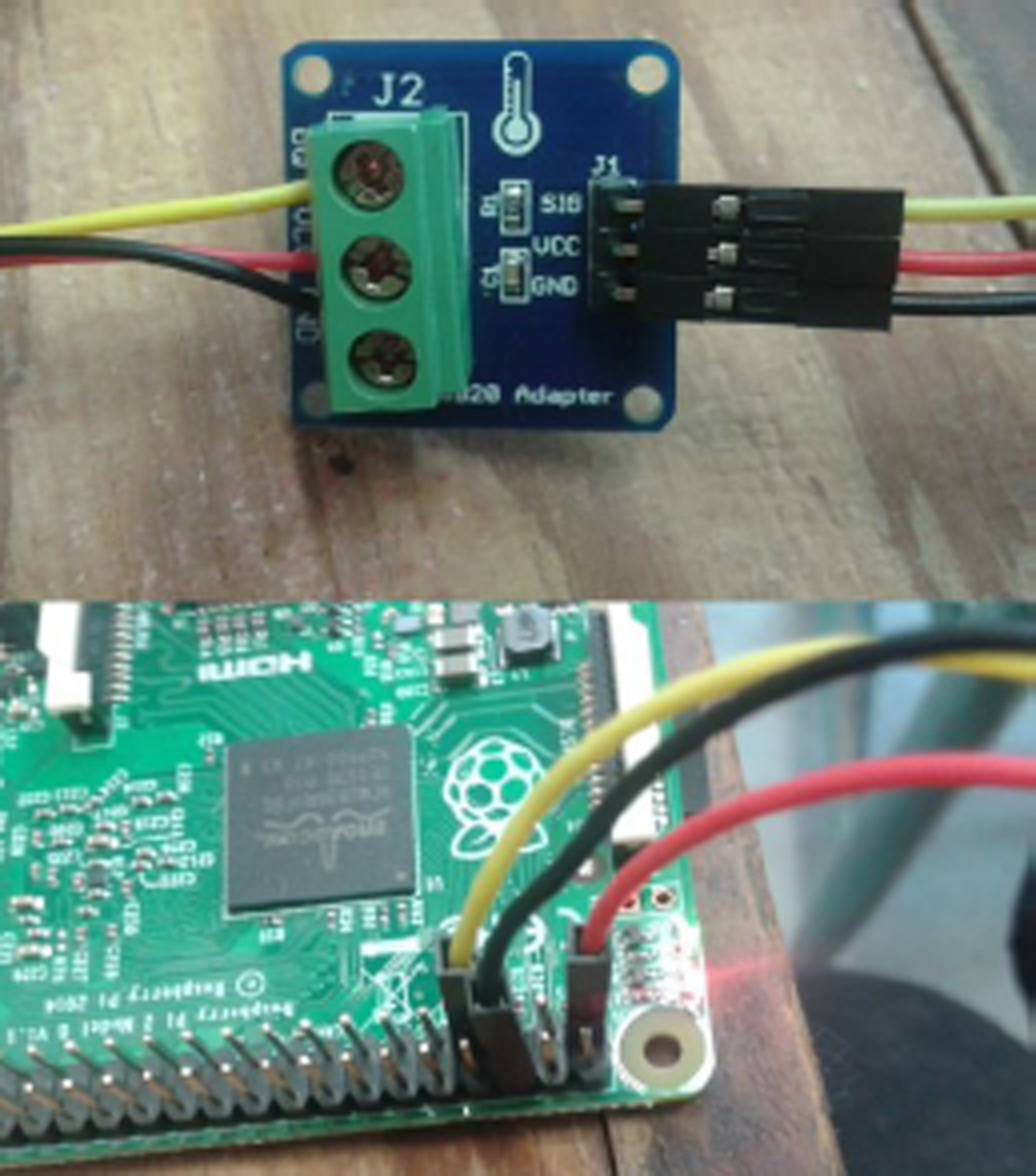
Shows the adaptor between the 1-wire digital thermometer and the Raspberry Pi, and how to connect the adaptor cables to the Raspberry Pi.

To compare the performance of the sensors, a TidbiT and a low-cost sensor were left in room temperature water, then moved to two flasks at successively higher temperatures for approximately 6 minutes each.

#### Hydrophone Tests

Audio from the DolphinEar DE-PRO Balanced Hydrophone was recorded via Pure Data onto a USB flash drive (16 bit recording, 0 dB gain, single .wav audio file per kayak trip). This hydrophone has a recording bandwidth of 10 Hz - 24 kHz with an approximately flat (±2 dB) response 20 Hz - 20 kHz (-6 dB at 10 Hz and 24 kHz).

All acoustic data processing was carried out in MATLAB (2016b, The Mathworks). Tones at frequencies 10 Hz, 60 Hz, 200 Hz, 1000 Hz (1 kHz), 10000 Hz (10 kHz) and 20000 Hz (20 kHz), with known voltages as found using an oscilloscope, were recorded through the entire system (Raspberry Pi, pre-amp and sound card). The recorded voltage was then divided by the input voltage for each frequency to calculate the correction factor required. The values were interpolated using a standard linear interpolation method in MATLAB to provide a correction factor per 1 Hz.

During initial tests, paddling noise was pervasive - therefore for the purposes of scientific data collection the sampling procedure was to collect unpolluted samples by pausing paddling. Test trips were made with two Sonic Kayak systems on the Penryn estuary in Cornwall in March 2017 on an incoming tide. Pausing paddling to collect unpolluted recordings was limited by the windy conditions at the time. As well as pausing periodically where possible we also paused when sounds of interest were heard through the onboard speakers, for example, boats.

The hydrophone sensitivity is not available from the supplier, therefore only relative dB levels are calculated. To calculate the sound levels per 1 Hz a Fast Fourier Transform (FFT) was applied sequentially to overlapping 1 second segments (Hann window, 50% overlap). This process converts the waveform signal in the time domain to the frequency domain. The calculated correction factor is applied during this process. The results are converted into dB using the equation:

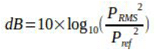

where *pRMS* is the result from the FFT process as root-mean square pressure, and *pref* is the reference pressure (the decibel scale is a logarithmic ratio scale). In underwater acoustics, the reference pressure is 1 μPa which is different to the reference pressure used for sound in air and they are therefore not directly comparable.

The root-mean-square Sound Pressure Level (SPLRMS) represents the mean broadband sound level for a given time period over a wide frequency range. Mean SPLRMS (1 s window, 50% overlap; 10 Hz - 20 kHz) were calculated for each period of paused paddling. For each 1 s window, an FFT process was first applied and the frequency-dependent correction factors applied. The result was converted back into the waveform and the SPL was calculated using the equation:

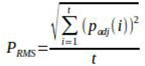

where *pRMS* is the root-mean-square pressure, *padj* is the corrected instantaneous pressure and *t* is the time (in number of datapoints). The mean SPLRMS was calculated for each period of paused paddling and then converted into dB using the same equation as above.

#### Electronics and Housing Design

The Sonic Kayak system is run by a Raspberry Pi 2 (£29). This is connected to a USB GPS dongle (GlobalSat BU-353-S4 USB GPS Receiver, £29), a 5V/3A battery (Anker Astro E5 16750mAh Portable Charger, £30), two powered speakers (Minirig, £120 each), a USB soundcard (CSL C-Media USB Mini Sound card, £10) leading to a pre-amp (Tascam iXZ, plug modified to allow connection to a standard microphone input, £32) leading to a hydrophone (DolphinEar DE-PRO Balanced Hydrophone, £280), two digital thermometers (DX Waterproof DS18B20 Temperature Sensor with Adapter Module for Arduino, £3 each), and a USB flash drive (Cruzer Glide 16Gb, £5) (Fig. 2).

**Fig 2.**
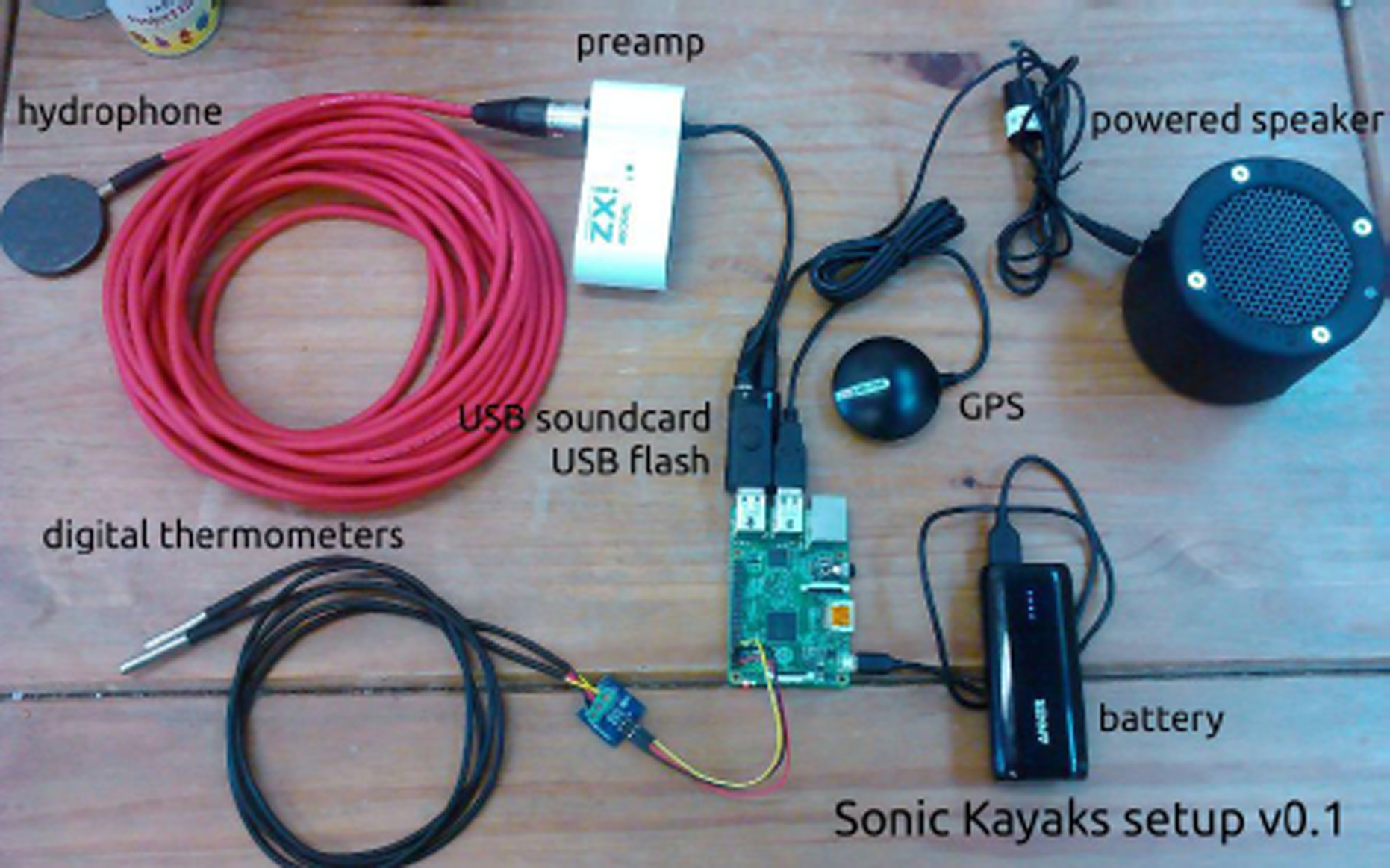
Hardware in the Sonic Kayaks.

The Raspberry Pi, GPS, battery and preamp were housed in a waterproof box, while the speakers were housed in customised cylindrical tupperware tubs. The box and speaker tubs were joined with flexible metal links (screw-in D-loops and metal clasps, easily found in any chandlery or good DIY shop). The box and speaker tubs were placed on top of the kayak at the front, and attached to the kayak using custom designed adjustable nylon webbing straps, running under the bottom on the kayak and attaching to the handles on either side of the kayak. The hydrophone was placed off the front end of the kayak (secured with cable ties) with a weight close to the sensor and a float approximately 1m along the cable to keep the sensor at a depth of 1m. The digital thermometers were placed over each side of the kayak, with a weight close to the sensor and a float approximately 60cm along the cable. Neoprene pads (1cm thick) were cut to fit under the box and speaker tubs to ensure a tight fit to different kayak models. In total, the material for fixing and waterproofing cost approximately £60. We designed and 3D printed gramophone-style horns to top each of the speakers (shape files available open-source here: https://github.com/sonicbikes/sonic-kayaks/blob/master/hardware/horn-print.stl, print costs £60 each via Shapeways.com). The complete system (Fig. 3) cost approximately £841 in materials, though depending on needs, it would be possible to substantially reduce this cost by omitting or replacing the hydrophone, speakers and/or printed horns with cheaper versions.

**Fig 3.**
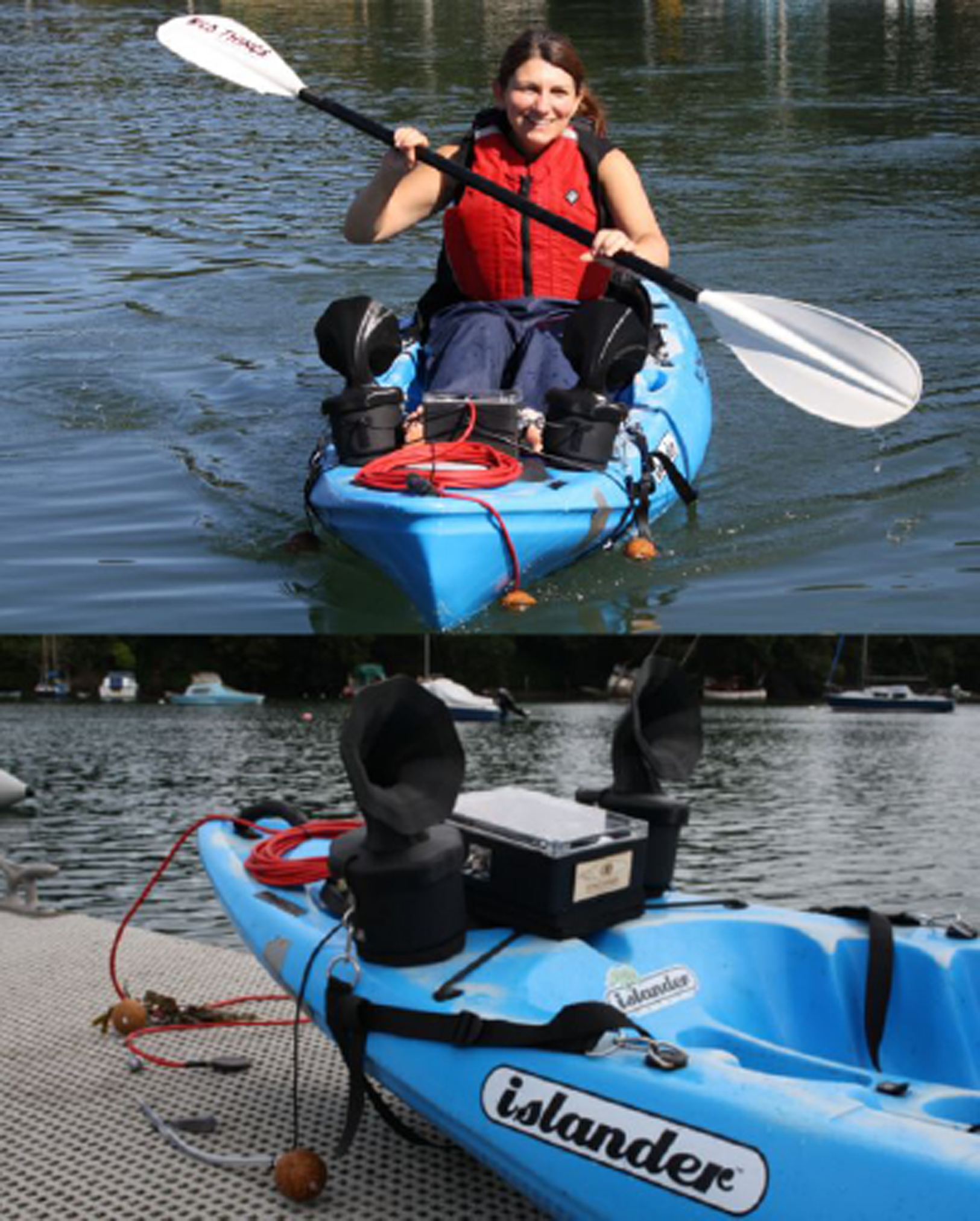
The Sonic Kayak system

### Sound Design

We used three layers of sound on the Sonic Kayaks - real-time sonification of the data obtained from digital thermometers, a live-feed from a hydrophone, and GPS-triggered sounds placed in different geographical locations. Below each layer of sound is addressed in turn.

#### Digital Thermometer Sonifications

Data from the two digital thermometers on each boat were sonified in real-time. As the water temperature goes up the sound pitch goes up, and vice-versa, with no sound when the temperature is stable. This was designed to provide easy-to-understand sonification of the underwater temperatures while not producing unnecessary noise that could detract from the peaceful nature of kayaking. The pitch is normalised to the running minimum and maximum the kayak has observed. This makes it much more sensitive, but it takes a few minutes to self calibrate at the start. Currently the pitch ranges from 70 to 970 Hz, with slight frequency modulation at 90 Hz to make the lower end more audible.

#### Hydrophones

A hydrophone was used to transduce live underwater sounds continuously. Even with a preamp the hydrophone needed to be boosted in Pure Data by 12 times to allow the paddler to hear detailed sounds.

#### GPS-Triggered Sound Zones

Geographical locations were mapped with sounds attributed to them, using custom software. The GPS on the kayaks detected when the boat was within one of these zones, triggering the sounds to play. For the British Science Festival, we mapped three bands of sound running out from the shore (Fig. 4), to allow for the tide changing dramatically over the course of the day (Swansea Bay has the longest tidal range in Europe). One zone contained poetry about climate change and the sea, another zone contained beats and music, and the third zone contained electronic voices with factual messages about climate change. Between each zone there were areas where no additional sounds were present, allowing the paddler to choose how they wanted to use the kayaks.

**Fig 4.**
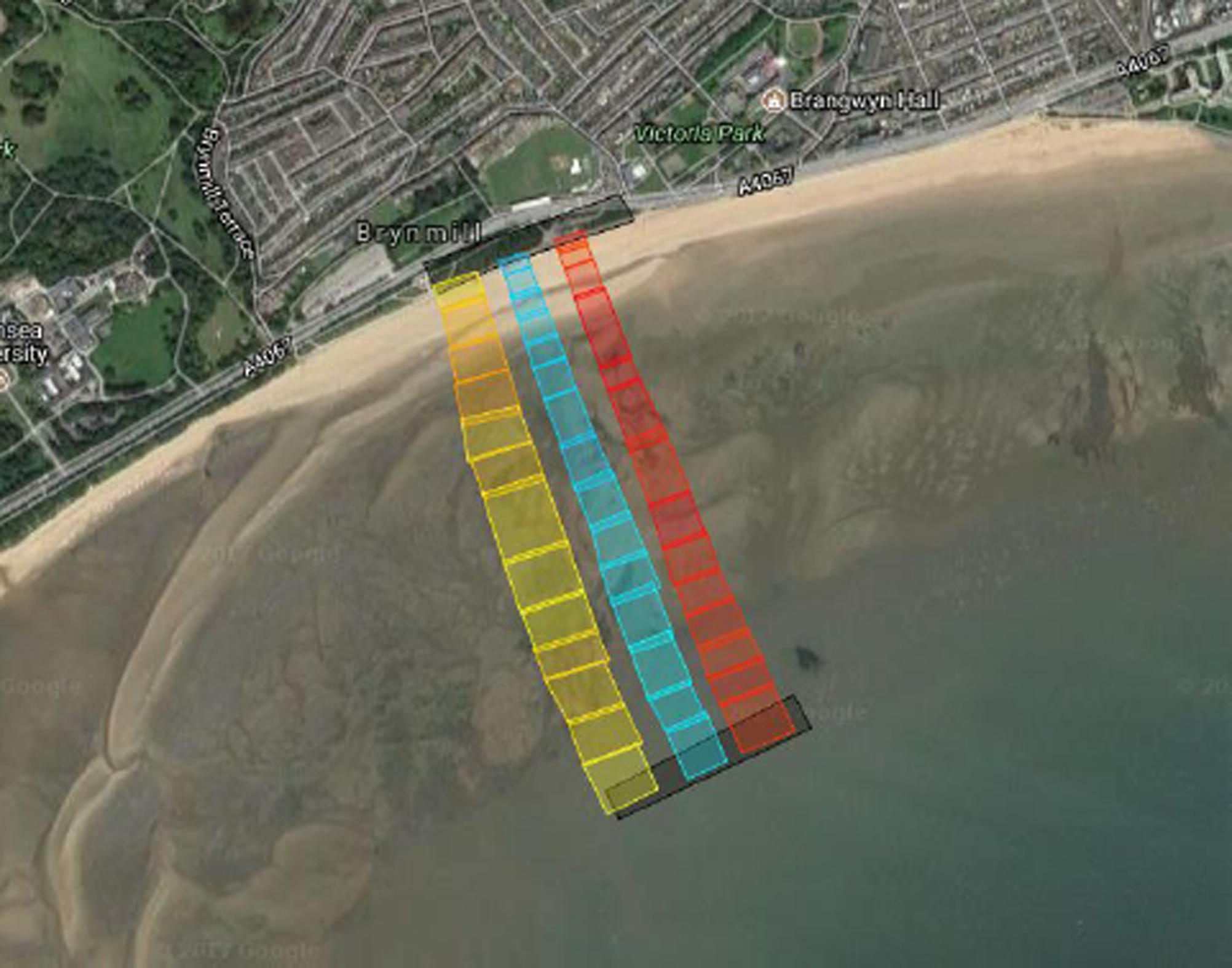
Sound zones as entered in the mapping software - each colour represents a different type of sound, and each small block refers to a different sound sample - when the kayak moves between each region the different samples are triggered, fading out and in as boundaries are crossed.

### Sonic Kayak Software

The software is built from three independent systems, one is written in Lua and is based on the original Sonic Bike system. This is responsible for reading the GPS device and creating events for when the kayak enters or leaves a GPS zone defined on a supplied map. This system also records the GPS location in a logfile every second. The temperature sensor driver is written in Python and reads the two digital thermometers using the 1-wire serial protocol on the Raspberry Pi General Purpose Input/Output (GPIO) port, logs the reading and creates a secondary calibrated value to give highly sensitive indications of rises or falls in temperature. These readings, along with the enter/leave zone events are passed to the sonification system via Open Sound Control messages (OSC). The sonification system is written in Pure Data, an open source visual programming language. Pure Data converts the thermometer data into sound - via sample playback in the case of zone events, or synthesising the sound directly from the data in the case of temperature changes. The sonification system also reads the hydrophone sound, amplifies it and passes it directly to the output - which is fed to the two speakers. Figure 5 provides a schematic of the different systems and how they feed into each other. The software running on the Sonic Kayaks is available open-source here: https://github.com/sonicbikes/sonic-kayaks

**Fig 5.**
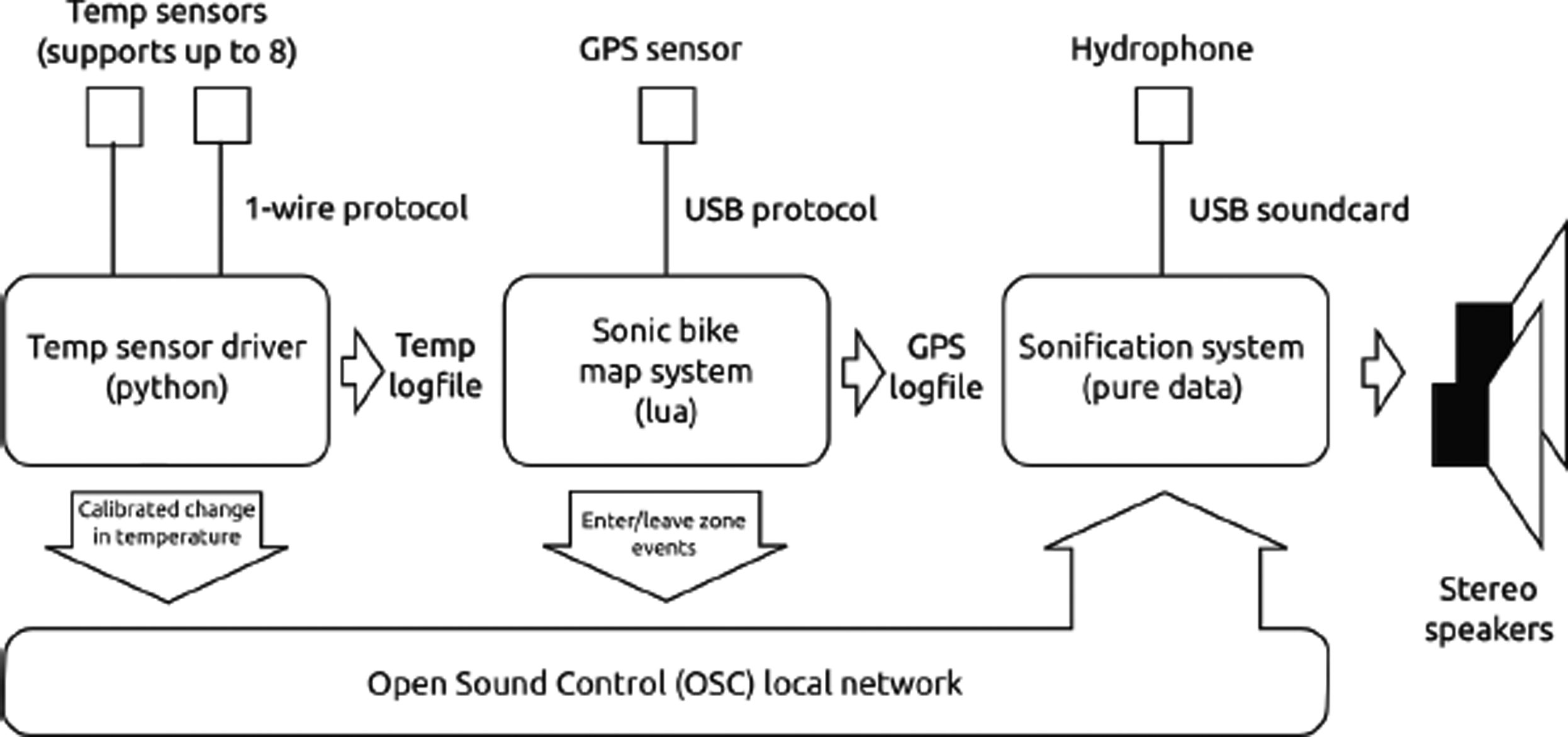
Sonic Kayak software architecture

### Citizen Collaborations

#### Development Hacklabs

Instead of developing this project as a small closed team, we launched open invitations for participation and collaboration via two hacklab events, to attract more diverse skills and allow co-creation from the outset and co-learning across disciplines. Prior to the first hacklab, we had not started the development process, as such a broad range of people and perspectives were involved from the very beginning. During these events participants were involved in designing and prototyping the system, including working with sound (recording from nature, and sounds generated through instruments and algorithms), developing the audience experience, testing sensors, coding and electronics.

#### Festival Event Implementation

We built two Sonic Kayaks for use in a two-day installation at the British Science Festival 2016 in Swansea, UK. The event was fully booked with 64 participants. We ran four 45 minute sessions each day, with eight participants per session - within each session there were two sonic kayakers and six additional kayakers. Each session started with a 10 minute safety briefing, allowing time for the equipment to be checked and adjusted between groups. We worked with the 360 Beach and Watersports Centre on Swansea Bay for the delivery of this event - they provided the kayaks, paddles and safety gear, gave the safety briefings and took the groups out each time. We used a pop-up tent and solar panel for storing, fixing and charging equipment, and provided waterproof comments books for the participants on their return. The British Science Association used their standard feedback forms for the event, including (i) rating the event overall as Excellent, Good, Average, Poor, or Terrible, (ii) how has this event affected your interest in science? - I’m much more interested, I’m a bit more interested, My level of interest is unchanged, I’m a bit less interested, or I’m much less interested, and (iii) asking participants for three words to describe the event. We uploaded the temperature data open-source immediately following the event. Hydrophone data was not recorded during this event as this feature was developed later.

## Results

### Sonic Kayak Hardware

#### Temperature Sensor Tests

The low-cost digital thermometers monitor the temperature every second, giving much greater resolution than the TidbiT which took ~6 minutes to reach the correct temperature - in our tests the low-cost sensor data shows the water in the flasks cooling over time, while the TidbiT is still slowly honing in on the temperature (Fig. 6).

**Fig 6.**
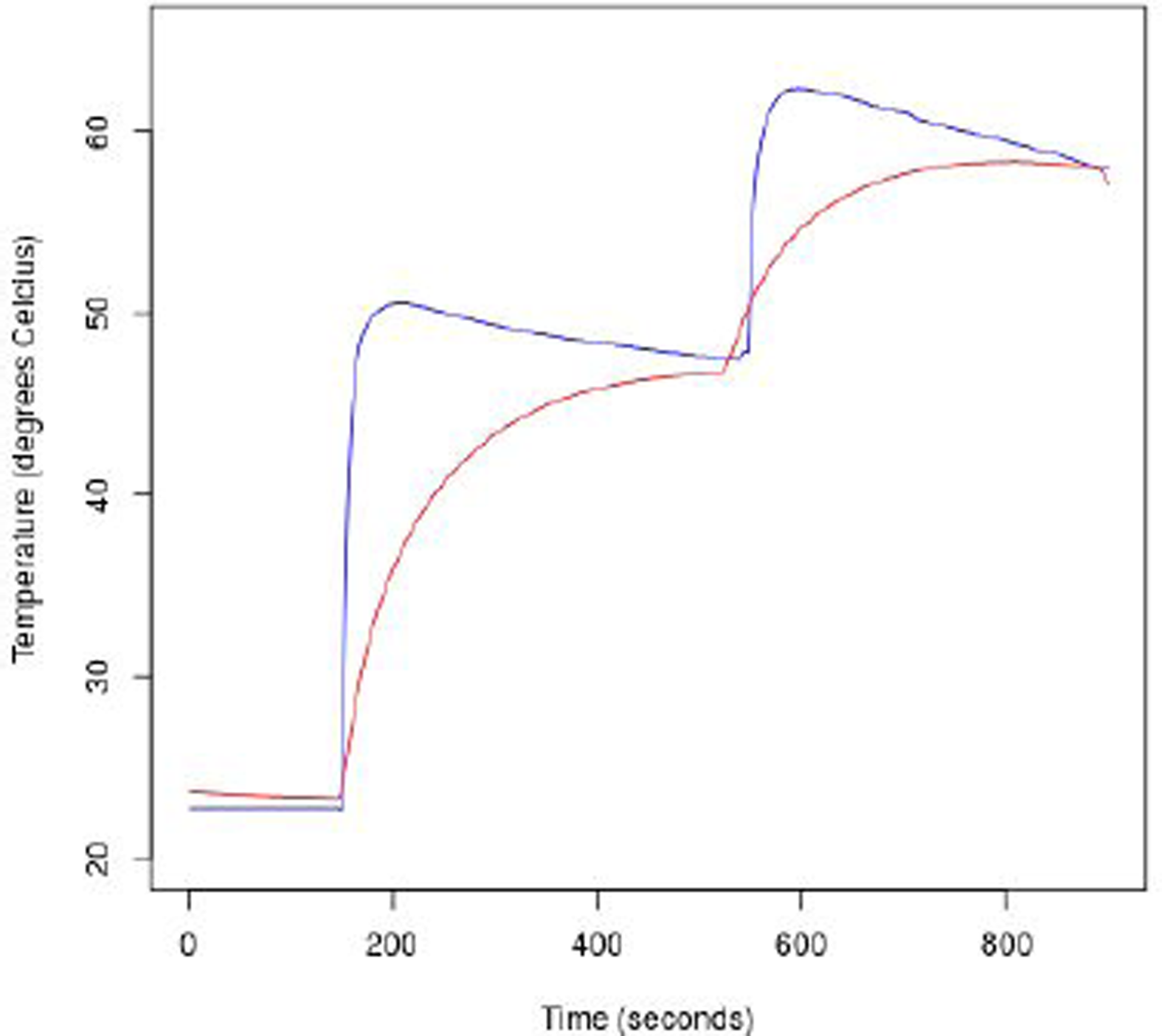
Temperature data obtained from a TidbiT sensor (red) and a low-cost digital thermometer (blue), starting in water at room temperature, and moving into two flasks of water at successively higher temperatures for ~6 minutes each.

The datasheets for indicate that the low-cost sensors are slightly less accurate than the TidbiT (TidbiT Range: −20 to 70°C, Accuracy: ± 0.21°C, Resolution: 0.02°C, Response time: 5 mins; Low-cost 1-wire digital thermometer Range: −55 to 125°C, Accuracy: ± 0.5°C, Resolution: 0.063°C, Response time: not stated), however our results indicate that the low-cost sensors provide better and faster data resolution, with an accuracy level equivalent to the TidbiT.

#### Hydrophone Tests

Figure 7 shows the path taken during the hydrophone tests, as monitored by the on-kayak GPS. Self-noise was present at several frequencies with peaks at 3,798 Hz, 4,138 Hz, 7,925 Hz, 8,679 Hz, 11,712 Hz, 12,809 Hz, and 20,679 Hz - this is thought to originate from the Raspberry Pi or the GPS after the elimination of all other components. Paddling produced a broad peak ~300 Hz and peaks at 500 Hz and ~2 kHz (Fig. 8). When paddling, the noise masks other sounds at similar frequencies.

**Fig 7.**
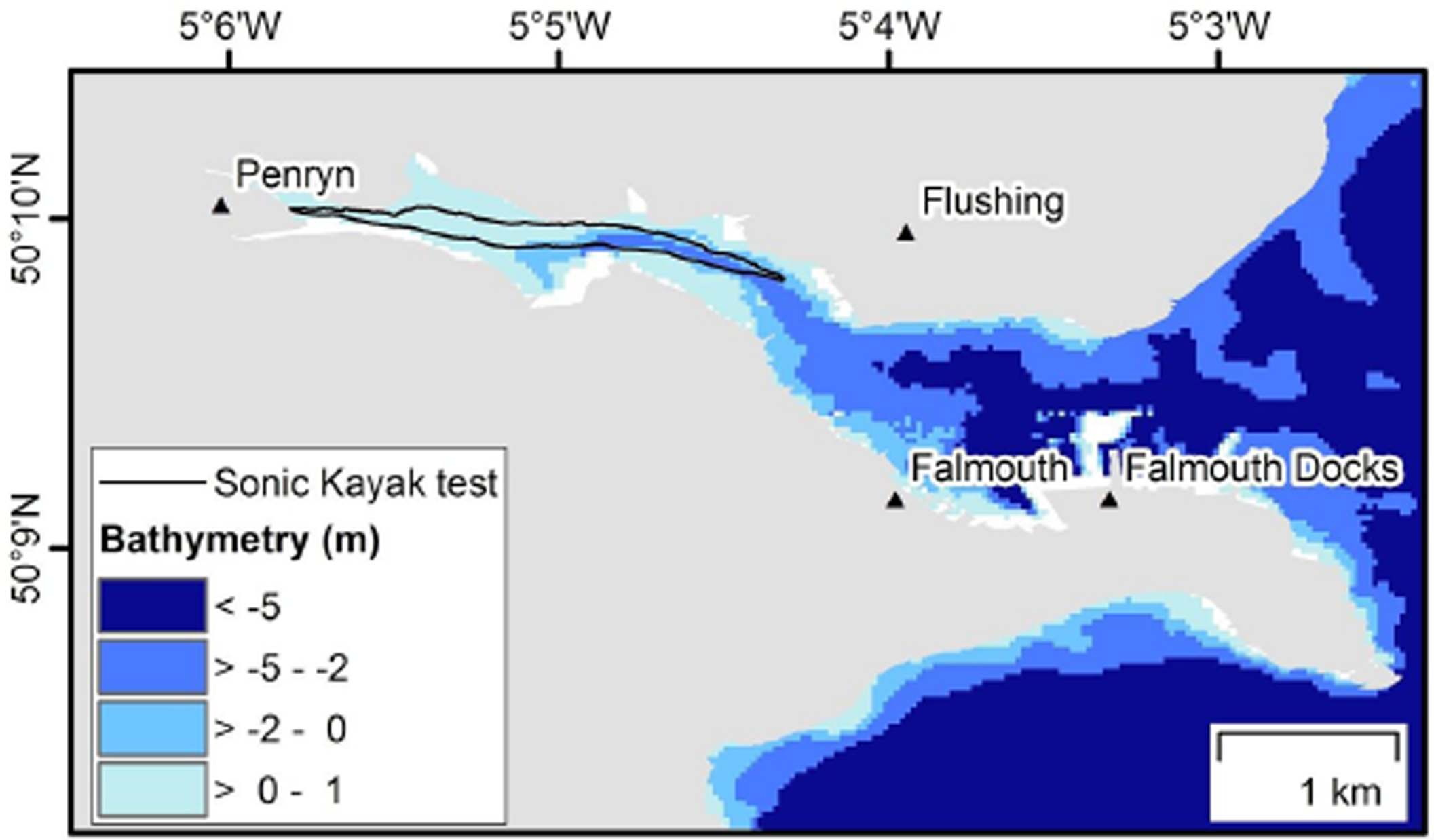
Map showing the Sonic Kayak test March 2017. The depth approximates to Lowest Astronomical Tide (LAT). Bathymetry data source: SeaZone HydroSpatial One, 1 arcsecond (~30 m) gridded bathymetry © Crown Copyright/SeaZone Solutions. All Rights Reserved. Licence No. 052006.001 31st July 2011. Available at: https://digimap.edina.ac.uk/ Accessed on: 12th April 2017.

**Fig 8.**
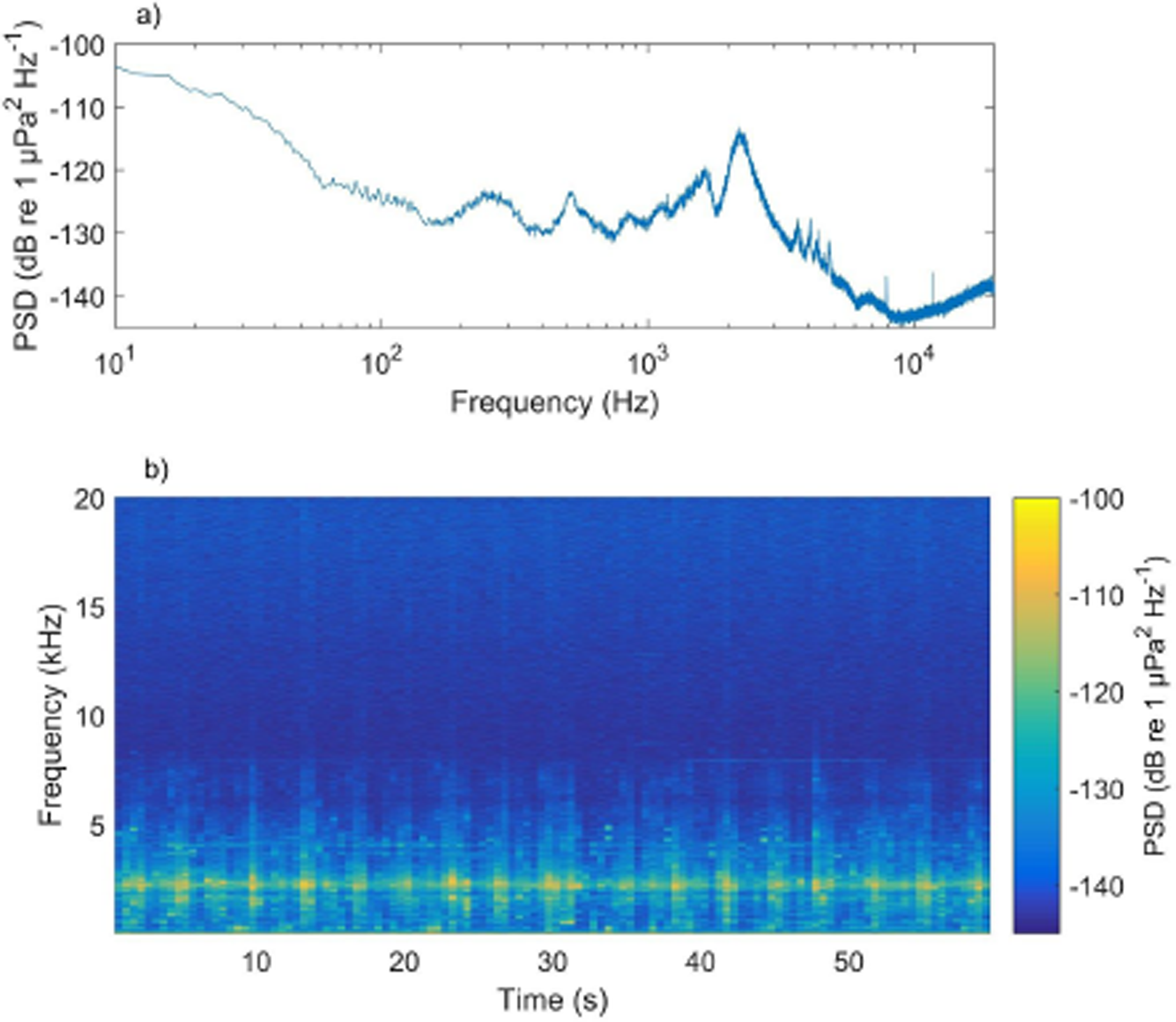
a) Mean power spectral density (relative dB) of the paddling noise given in (b), giving the average relative sound levels per 1 Hz for the entire duration presented in (b). b) Spectrogram of a sample of paddling noise displaying the sound levels per 1 Hz per 1 s (50% overlap; time interval is 0.5 s).

Periodically and when sounds of interest were heard through the speakers, paddling was paused to record unpolluted samples. For each of these samples, the mean broadband sound level was calculated for the frequency range 10 Hz - 20 kHz (SPLRMS re 1 μPa). These levels varied considerably throughout the Sonic Kayak sound test by 14.6 dB (Fig. 9). The loudest SPLRMS (identified in red in Fig. 9) occurred near to Falmouth Marina and consisted of peaks at specific frequencies from 200 Hz - 10 kHz (Fig. 10).

**Fig 9.**
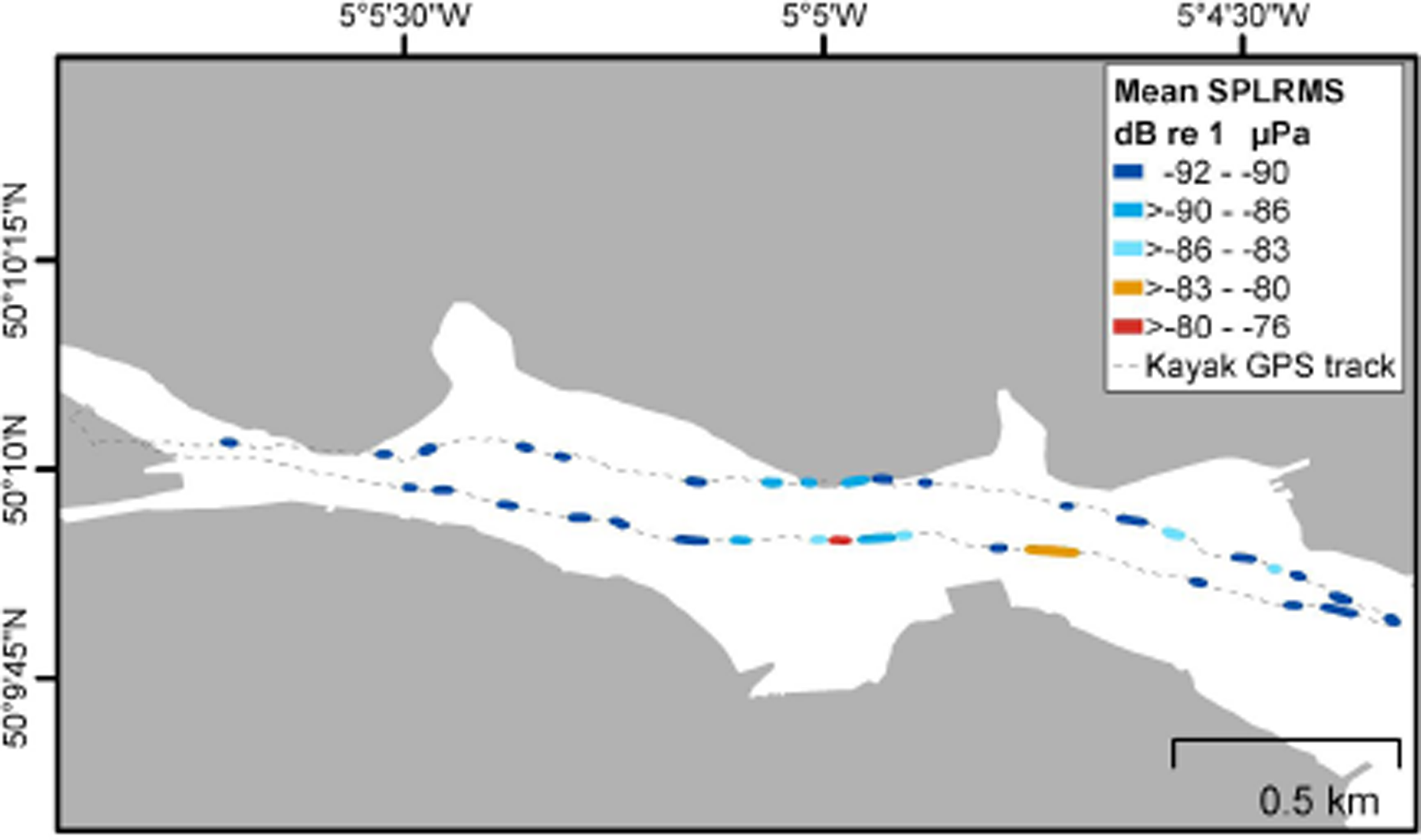
Mean relative SPLRMS dB re 1 μPa (10 Hz - 20 kHz) for each section of paused paddling. The sections are displayed with a 10 m buffer for ease of visualisation.

**Fig 10.**
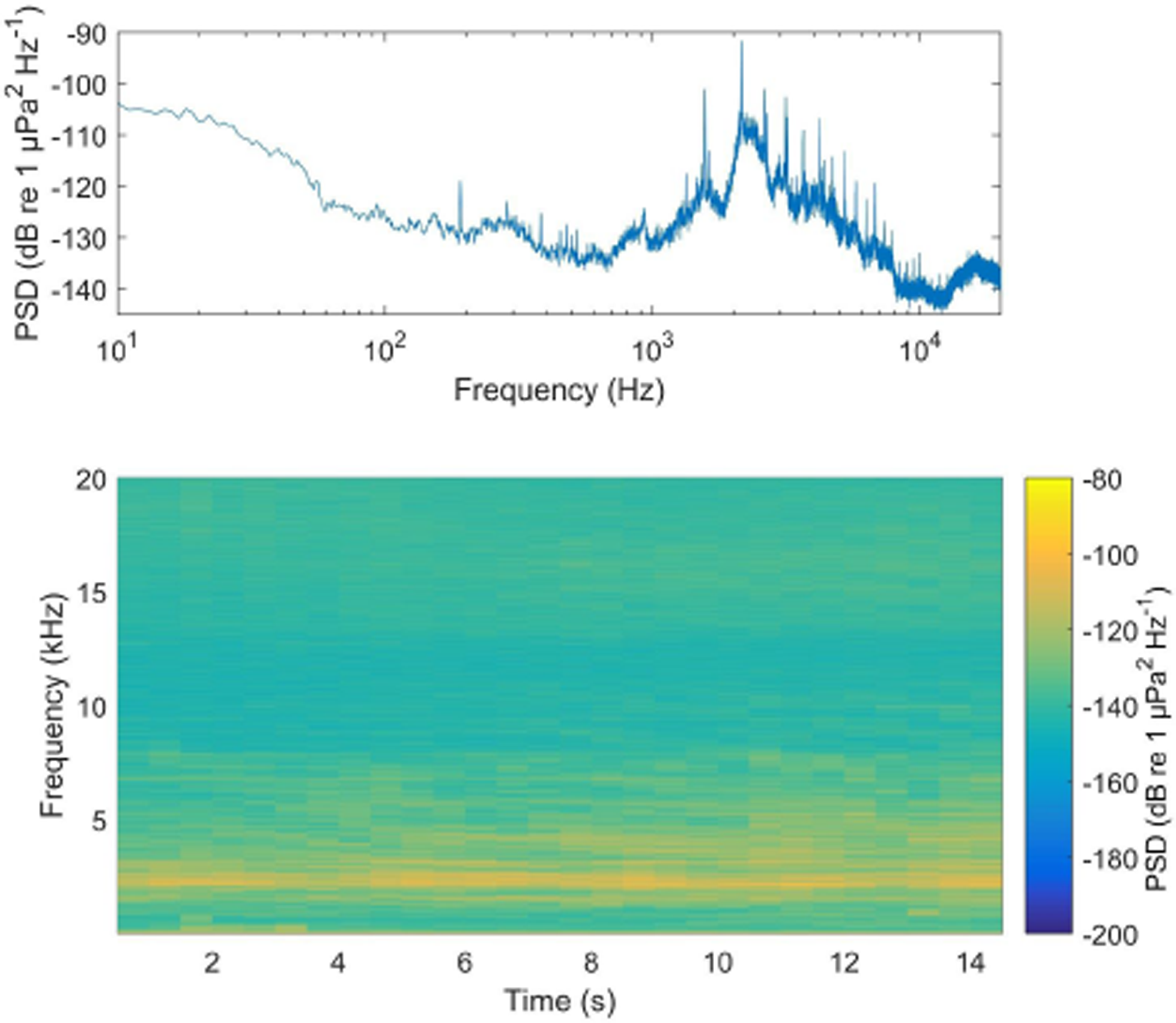
Mean power spectral density (relative dB) of the loudest mean SPLRMS identified giving the average relative sound levels per 1 Hz for the entire duration presented in (b). b) Spectrogram of the loudest mean SPLRMS identified displaying the sound levels per 1 Hz per 1 s (50% overlap; time interval is 0.5 s).

#### Electronics and Housing Design

The electronics and housing design worked well throughout four hours of continual use on the first day of the British Science Festival. However, we did notice some corrosion from salt water on metal components of the Raspberry Pi and speakers that had voltage differentials. In addition, towards the end of the four hours, there was condensation inside the main electronics box, caused by the components heating up inside, surrounded by a colder environment outside the box. On the second day, both kayaks were capsized for several minutes due to difficult waves, and the session was called off. Remarkably, one set of Sonic Kayak hardware remained fully functional with the exception of one speaker - for the other set the waterlogging was terminal.

### Sound Design

Recordings from the Sonic Kayaks are available here: https://archive.org/details/sonickayak-recordings. A similar shorter clip was played as part of a 30 minute interview about the Sonic Kayaks on the Cerys Matthews show on BBC Radio 6 Music 28 August 2016 http://www.bbc.co.uk/programmes/p0464c3r, demonstrating the wide reach attained by taking this transdisciplinary approach.

#### Digital thermometer sonifications

Originally we designed the temperature sonification to be continuous. It proved important that no sound was produced when the temperature was stable as during testing, users remarked on wanting more quiet time while out on the water. This also served to keep kayakers aware of their above-water surroundings, which is advantageous from a safety perspective.

#### Hydrophones

During testing in an estuary setting, the hydrophone gave insight into man-made noise pollution particularly from boat motors, and noises from wildlife and the motion of the water. Boat sounds are very loud through the hydrophone - it is possible to hear engines over long distances, which is useful in expanding your senses if the boats are out of sight. This is because sound travels five times faster underwater than it does in air at ~1500 ms-1, and so propagates much further. We also heard clicking sounds from underwater species, likely snapping shrimp, chains clinking, background ship noise, flow noise, wind and waves. In Swansea Bay the environment was much more homogenous underwater (sand) and noisy above water (waves) – the hydrophone contributed very little in this setting, so having the additional GPS triggered sounds proved an important addition. While testing in the relatively quiet estuarine setting, the system could clearly be heard from onlookers on land - in this case, as with the Sonic Bike system that came before, the paddlers inherently become performers and passers-by become an audience.

#### GPS-triggered Sound Zones

During testing it became apparent that there were divergent views on this additional layer of sound. Some users preferred more quiet and only to have the temperature sonification and hydrophone sounds, while others wanted additional sounds to make the experience more interesting. The final design for the British Science Festival implemented bands of sound allowing the paddler to choose their experience, as shown in the methods section.

### Citizen Collaborations

#### Development Hacklabs

Participants came with backgrounds in climate change, algal research, fisheries conservation, drone operation, e-textiles, art workshop provision, and computer games. We found it extremely beneficial to have such broad perspectives from the very start of the development process. Much of the software, build design and sound design used in the final version were developed during these sessions or designed based on feedback from these sessions. We recommend this approach for any citizen science project, or indeed any research project intended to be of use to others.

#### Festival Event Implementation

A video explainer from the British Science Festival event is available here, showing the system, how it works, and some of the sounds https://vimeo.com/184935959. Our main recommendation for others wanting to do similar interventions is to ensure that a watersports specialist is actively involved in the project. For the festival, we worked with the 360 Beach and Watersports Centre, who ensured that the basic health and safety issues associated with watersports were met.

Participant feedback was collected by the British Science Association and 21 feedback forms were completed. When rating the event in general, 18 people rated it as ‘excellent’, and two rated it as ‘good’ - one person did not provide a rating. When asked if the event affected their interest in science, five people said they were ‘much more interested’, nine said they were ‘a bit more interested’, three said their ‘level of interest was unchanged’, and three did not provide a rating. A total of 17 people provided three words to describe the event, and two people provided two words - these are available as supplementary material S1a. Feedback gathered from our comments books and talking to participants informally indicated that, after taking the kayaks out, many of the participants understood the climate change aspect to the project despite no specific introduction or discussion of the subject. Two participants were visually impaired and highlighted the possibilities for such a system for auditory explorations of different environments. Some participants had come simply for the opportunity to use a kayak for free. Written comments are available in supplementary material S1b.

Figure 11 shows the data collected by the festival participants on day one, mapped over a satellite image. Each map is from one of the two Sonic Kayaks, who took roughly the same routes - the similarity between the data indicates that it is replicable (Kayak 1: Min temp = 17.687°C, Max temp = 25.687°C, Mean temp = 19.712°C, n = 14,431 records; Kayak 2: Min temp = 17.812°C, Max temp = 23.125°C, Mean temp = 19.792°C, n = 13,462 records). On each map the four consecutive trips from day one have been overlaid. It is possible to see where the boats were taken out of the water between trips (the yellow regions where the temperature rapidly goes up), and how with successive participant groups as the tide moves out, the water temperature increases – showing how the kayaks could be used to detect spatial and temporal microclimatic changes. The raw data is available here:https://github.com/sonicbikes/sonic-kayaks/tree/master/sensors/viz/archive

**Fig 11.**
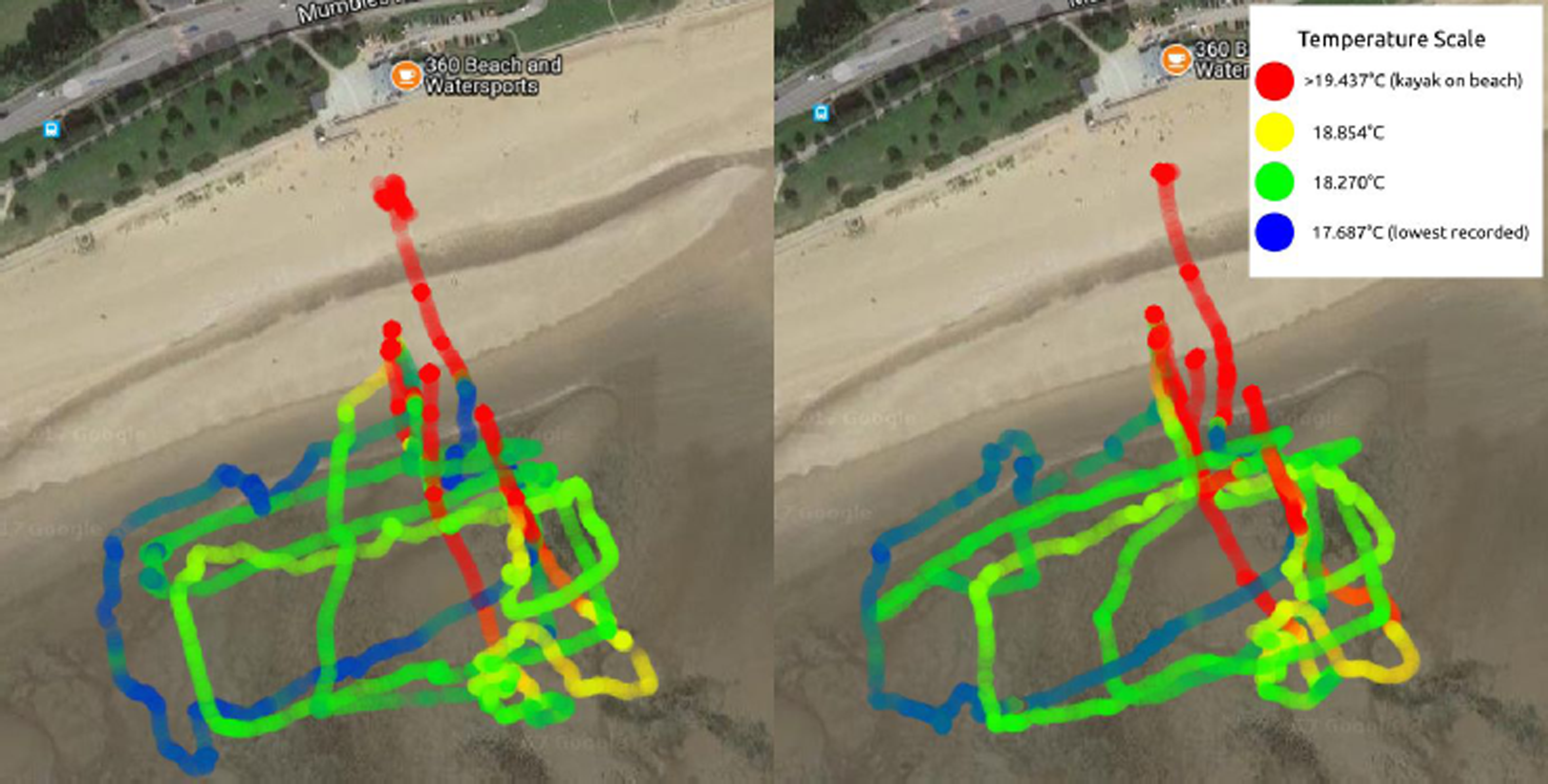
Maps showing the temperature data collected by the two Sonic Kayaks at the British Science Festival - each map shows the data from one of the kayak systems, with the four consecutive trips overlaid. Blue indicates lower temperatures, while red indicates higher temperatures. This demonstrates the possibilities for obtaining very fine-scale maps of temperature data over space and time.

## Discussion

The Sonic Kayak became a musical instrument, a portable stage, a game player, a doorway to listening and looking that creates good health, and a tool with which to collaboratively monitor our changing environment. The launch event at the British Science Festival was fully booked and received good press coverage (BBC Radio 6 Music Cerys Matthews show, BBC Radio Wales, BBC World Service, local print media), indicating widespread appeal from the combination of sound art, climate and marine science, and sport. The system has proven potential as a tool for researchers in the fields of underwater noise and climate science.

The use of similar low cost open-hardware equipment is becoming increasingly popular for marine research. Researchers have developed a static underwater recording device based on a hydrophone and Raspberry Pi for deployment at fixed locations around the Brazilian coast to measure the impact of industrial developments on the marine ecosystem (Caldas-Morgan et al. 2015). An anchored inflatable kayak has also previously been used to deploy a hydrophone at selected sampling locations on a shallow coral reef with the intention of reducing wave slap noise on the hull of a normal research boat (Nedelec *et al*. 2015). The maximum depth in this area was 0.6 - 4.5m highlighting the benefits of using a kayak in delicate and vulnerable shallow water environments where minimising disturbance is a goal. The potential for surfers to collect environmental data has been demonstrated by Brewin et al. (2015), who fitted TidbiT sensors to surfboard leashes, and GPS packs to the surfers - they estimated that each year, 40 million measurements on environmental indicators could be generated around the UK coastline by surfers. This idea is currently being turned into a commercial venture by smartfin http://smartfin.org/ who are making a surfboard fin with integrated sensors for salinity, temperature, location, wave characteristics and pH. The Sonic Kayak system takes this approach further by providing feedback to the user, making it more likely that people will enjoy using it and actively seek out the experience - this aspect is frequently not considered, yet it is critical for encouraging uptake and retaining interest.

### Applications for Underwater Sound Research

It is widely recognised that anthropogenic underwater noise has increased, particularly in the northern hemisphere. This has predominantly been attributed to commercial shipping (Andrew *et al*. 2002; McDonald *et al*. 2006; McDonald *et al*. 2008). In addition to chronic increases in average sound levels, impulsive sources also affect underwater sound levels such as seismic surveys, sonar and construction (Hildebrand *et al*. 2009; Thode *et al*. 2010; Estabrook *et al*. 2016). Many marine species use underwater sound for communication, orientation and hunting (Versluis *et al*., 2000; Verfuß *et al*., 2005; Janik 2009; Radford *et al*., 2014). They are therefore at risk of adverse impacts from anthropogenic noise. This has been documented for a wide range of marine species. For example, shipping noise has been found to affect the foraging behaviour of humpback whales (Blair *et al*. 2016) and fish species (Magnhagen et al., 2017), alter the immune response of the European spiny lobster with potential increased risk of infection (Celi *et al*. 2015) and reduce the ability of fish (Lusitanian toadfish) to detect acoustic signals in their environment (masking; Vasconcelos *et al*., 2007). In extreme cases, anthropogenic underwater noise can cause mortality as occurred in Falmouth Bay, UK in 2008 where naval activity was thought to be responsible for causing the mass stranding and deaths of common dolphins (Jepson *et al*., 2013).

Knowledge of exposure to anthropogenic noise is required to identify potential consequences for marine species and develop management practices. However, underwater sound can be highly variable, particularly in shallow, coastal environments (Mercado *et al*., 1999). In these environments, factors affecting sound propagation vary throughout the water column, spatially throughout an area and/or temporally such as bathymetry and temperature (Dosso and Chapman 1987; Mercado and Frazer 1999; Hazelwood & Connelly 2005). Furthermore, there are a wide variety of sound sources including natural sources such as wind and surf noise, biological sounds such as snapping shrimp and anthropogenic sources including recreational and commercial vessel activity (Haxel *et al*., 2013; Buscaino *et al*., 2016; Garrett *et al*., 2016). Therefore, methods of collecting underwater sound data to capture some of this variability are required. The Sonic Kayak system could be used for capturing this spatial variability.

In addition to collecting data for use in anthropogenic noise research, the Sonic Kayak system has potential in mapping biological noise and associated biodiversity as a non-destructive sampling method. A similar approach was used in a complex estuarine environment where passive acoustic monitoring was used to locate fish spawning aggregations and to relate habitat characteristics (seagrass coverage) with the fish vocalisations, potentially informing management (Ricci et al. 2017).

Using a kayak for underwater sound data collection has several advantages compared to traditional methods of drifting, towed or static autonomous recorders: (i) It is possible to achieve more fine scale and greater spatial coverage as compared to a single fixed recorder. In our test, we covered a distance of >2 km of the Penryn estuary and captured a total variability of nearly 15 dB with the SPLRMS changing by a maximum of ~10 dB over <100 m. (ii) Sampling can be more carefully directed than is possible with free-drifting sound recorders (iii) The Sonic Kayak system is low cost as compared with an array of fixed standard recorders. (iv) Kayaks cause minimal disturbance in comparison to a typical research boat, and can reach shallow areas more easily with less impact. (v) There is also the additional benefit of engagement with a greater number of people, connecting people with the underwater environment that they can’t typically see.

One potential local application is the monitoring of mearl. Falmouth Bay (and the Helford) are designated a Special Area of Conservation (SAC), of particular importance are the maerl beds (JNCC 2016). Maerl is a slow-growing calcareous algae, and an important habitat for many marine species. The Sonic Kayak system could be used to record samples of underwater sounds over both dead and alive maerl beds to assist with assessments of biodiversity and future management. A similar approach has been utilised in other northeastern Atlantic maerl beds, where acoustic recordings were used to compare the ecosystem health of areas with different levels of fishing activity (Coquereau *et al*., 2017).

Disadvantages of this system for research use include the necessity to stop paddling for recording data due to the paddle noise, the unlikeliness of gathering data overnight and potentially dawn and dusk where biological noise is greatest (Radford *et al*., 2008a; Radford *et al*., 2008b; Ghazali *et al*., 2013; Ricci *et al*., 2017), and the possibility for limiting weather conditions. The Sonic Kayak system may be best used complementary to traditional fixed systems. There is also great potential to use it in combination with additional data collection. For example, Sonic Kayaks could be integrated into the “Paddle into Science” citizen science programme with the Community Seagrass Initiative, where cameras with associated GPS loggers are towed underwater from kayaks to gather data specifically for seagrass mapping (CSI 2017).

Depending on the application of the system, more research may be required to identify and remove the source of electrical noise. This may require replacing one or more of the components - if it is the Raspberry Pi then a new base computer itself would be required. Knowledge of the hydrophone sensitivity is also required for the calculation of absolute sound levels, and so this would be essential for comparison with recordings taken using different equipment.

### Applications for Climate and Ecosystem Research

Sea temperatures are typically measured either using temperature sensors like the TidBit attached to buoys, ships or shorelines, or via satellite observations which measure variations in the amplitude of electromagnetic radiation wavelengths caused by temperature change typically at spatial resolutions of a number of kilometers (e.g. Nasa’s Aqua satellite https://directory.eoportal.org/web/eoportal/satellite-missions/a/aqua).

Fine-scale spatial and temporal temperature data are not commonly obtained, but are critical for understanding coastal ecosystems which change over very small distances and throughout the day. Coastal zones include a number of critical ecosystems and account for 90% of marine fisheries catches - estuarine ecosystems in particular provide key habitat for fish and invertebrate breeding, foraging and predator avoidance, and are highly vulnerable to climate change and drought causing changes to freshwater flow (Barbier 2017, Williams et al. 2017).

The ability to obtain fine-scale temperature maps, both spatially and temporally, would be advantageous for the study of the current and future impacts of climate change on estuarine fish breeding habitats. Indeed, such technology could also be applied to bettering our understanding of why algal blooms occur in specific areas at certain times (Hallegraeff 2010, Paerl & Huisman 2008), monitoring the impact of commercial/residential outflows or harbours on ecosystem microclimates (Bamber 1995), or the identification of suitable locations for shellfish and fish conservation, restocking or farming (Bromley et al. 2016, Ellis et al. 2015, Searcy-Bernal 2013) for example, and could equally be applied to river and lake ecosystems as well as coastal. Additionally, the acoustic recording capabilities mentioned earlier could be used to monitor such conservation implications.

The digital thermometers tested offer a resolution and replicability that is suitable for research use. The only modification necessary, depending on the data requirements, might be to fix the depth of the thermometers to ensure that this is kept constant. Currently the sensors move higher in the water column the faster the kayak is moving and potentially during turns, and so the depth at which the temperature is recorded is not consistent. Similarly, it would be possible to fix the depth higher or lower in the water column depending on what data was required, or to obtain a range of depth temperatures from a single location by moving the sensor manually.

### Additional Suggested Modifications

When the system was trialled in Swansea Bay for the British Science Festival, we experienced fast corrosion from salt water on metal components of the Raspberry Pi and speakers that had voltage differentials – as such, the system would benefit from further waterproofing. The current design is suitable for lake, river and estuarine use in dry weather, but is not suitable for use where there are waves or heavy rainfall, nor is it robust enough for long-term implementation. Options for waterproofing could include using resin to cover the electronics, and hermetically sealing the cable ports. In addition, towards the end of four hours continual use, there was condensation inside the main electronics box, caused by the components heating up inside, surrounded by a colder environment outside the box. Improving the waterproofing would no doubt help by reducing the flow of humid air into the boxes.

Additional sensors can easily be added to the existing system, for example instructions for low-cost open-source turbidity sensors are available and would enable fine-scale monitoring of pollution in estuarine waters from farm runoff (Kelley et al. 2014). During our second development hacklab, one participant added a sonar device (Lowrance elite-5 chirp) to the kayaks, together with a GoPro underwater camera to check whether the sonar information matched what was going on underwater. The participant (Dr. Chris Yesson, Institute of Zoology, London) is developing the use of sonar on kayaks to map kelp, a critical habitat for fish breeding. The system is fully open source allowing anyone to make their own version with any required modifications.

The system provides opportunities for citizen-led data collection. In future, a live mapping system addition would be beneficial, updating in real-time and showing where data gaps were to guide people towards particular locations. In terms of citizen-science data collection, GPS-triggered sound zones could bias where users spend time, and so bias the data collected. However, sound zones could also be used as incentives to move to specific areas, or as ‘rewards’ for entering an area that few people have gathered data from.

## Conclusions

Testing indicates that the Sonic Kayak system has potential as low-cost research equipment, immediately in the fields of climate change and marine noise, but also potentially in other fields related to water monitoring. The open hacklab approach proved highly valuable, bringing broad perspectives into the development process from the very start, helping to ensure that the Sonic Kayak met its dual purpose both for citizen science and sound art. We strongly recommend this approach for any research or citizen science project. Implementing projects that span disciplines brings difficulties – collaborations require patience and humility in order to produce a result that is considered high-level according to the vastly different expected outcomes of those involved. The payoffs are great however, both fuelling the inspiration of those involved and producing an end result that is more than the sum of its parts. Combining sound art and sonification with citizen science means that paddlers can use sound to explore and investigate the marine environment, and encourage further use of the system as it is an experience in its own right. We found that several participants at the British Science Festival had come simply for a chance to use a kayak, highlighting the additional benefit of the sports angle to the project for attracting participants who might otherwise not be interested. The sound element also opens unusual opportunities for those with visual impairments, further broadening the range of people who can benefit from such systems. It seems fair to conclude that projects which utilise a wide disciplinary approach will be of interest to a broader range of people. This can only be beneficial for citizen science.

## Acknowledgements

Funding for transdisciplinary projects is not straightforward – to launch this project we received funds covering the costs of equipment from the FEAST Cornwall (Arts Council England in partnership with Cornwall Council) and the British Science Association – both are forward thinking organisations who are not afraid to merge disciplines or to take risks on non-standard projects. We thank all those who came to the development hacklabs and who participated in the event during the British Science Festival 2016. We also thank the 360 Beach and Watersports Centre in Swansea for their fantastic support during the British Science Festival. Finally we thank Ivvet Modinou from the British Science Association team for taking a leap of faith and supporting this project from its conception.

